# Distributional invariance and proportional scaling in axonal conduction

**DOI:** 10.1101/2025.10.24.684120

**Authors:** Laurie D. Cohen, Shimon Marom

## Abstract

Conduction velocity along axons reflects geometric and biophysical influences whose joint statistical organization remains largely uncharacterized. Using high-resolution time-of-arrival measurements along hundreds of identified axonal branches in vitro, we quantified how propagation speed changes along trajectories. The ratio between terminal and initial velocities, *ρ* = *v*_end_*/v*_start_, follows a right-skewed distribution whose shape remains invariant across branch lengths, positions within neurons, and hierarchical aggregation levels. Local conduction profiles reveal a predominantly progressive deceleration along branches. These observations indicate a simple and robust statistical organization of slowdown, suggesting that proportional modulation of propagation speed is a consistent feature of axonal signaling in structurally variable substrates.

---

Several studies of the biophysical mechanisms underlying membrane excitability have converged toward a view of the excitable membrane as a self-organizing system whose kinetics exhibit scaling relations and functional resilience across wide ranges of structural variability [1–13]. These analyses, based on single-protein and whole-cell observations, suggest that reliable excitability arises from the collective kinetics of interacting conductances rather than from fixed molecular architecture. Here we ask whether such invariant properties extend to spatially distributed settings; conduction velocity along axons providing a simple test case. We examine this question at the level of spatially extended single neurons, providing a phenomenological account of conduction slowdown based on its statistical regularities.

Conduction velocity is a fundamental property of excitable media. In axons, it depends on geometry, membrane protein density, and multiple state-dependent processes that generate local variations in propagation speed [13–23]. Such variations can yield distinct macroscopic outcomes: if structural and kinetic factors balance geometric and branching effects, velocity remains invariant; if they do not, conduction may accelerate or slow along the trajectory. It may also vary idiosyncratically, reflecting unpredictable local contingencies.

We report that conduction slows down systematically along axonal trajectories. The slowdown is not idiosyncratic; rather it follows a statistical regularity: the distribution of the ratio between end-to-start velocities is right-skewed, and its shape remains stable across branch lengths and positions within neurons. These empirical regularities are compatible with multiplicative influences, extending current analyses of specific determinants toward a broader characterization of the stochastic processes that govern axonal propagation. Proportional scaling laws of this sort are familiar in transport through disordered or heterogeneous media, where local fluctuations compound multiplicatively to yield log-normal behavior [24–27].

All experiments were performed in *ex vivo* developing cortical networks [9, 28, 29]. Recordings were obtained with CMOS-based high-density multielectrode arrays (HD-MEAs; *MaxWell Biosystems* AG), a non-invasive platform enabling reconstruction of axonal trajectories from electrical activity [17, 30–32]. Axonal branches were identified with the *AxonTracking* module of the *MaxLab Live* software, which links sequential electrodes exhibiting time-shifted spike-triggered averages consistent with continuous extracellular conduction along axonal processes (Figure 1).

**FIG. 1.**
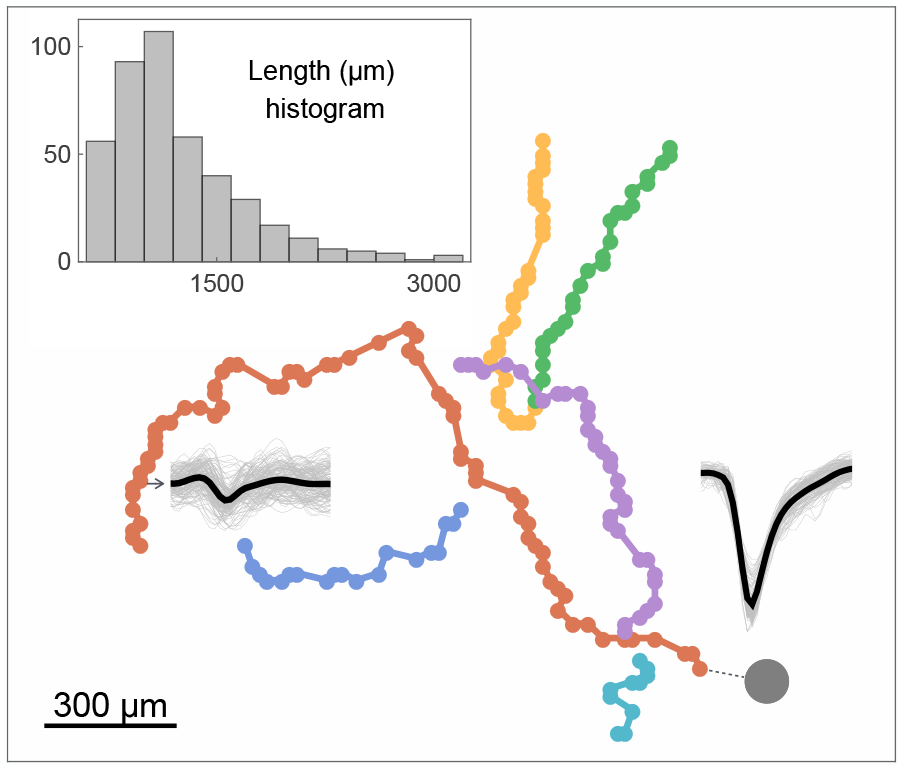
Axonal trajectories were identified using the *AxonTracking* module of *MaxLab Live*. The gray circle marks the site of somatic spike initiation, and the dashed line indicates the direction of propagation. Colored dots mark electrode positions underlying reconstructed paths. Representative voltage traces (thin lines, single events; thick lines, averages; trace duration ∼1.5 ms) illustrate a somatic spike followed by a spike in one distal axonal electrode ∼ 3.9 ms later. Scales: mean somatic spike amplitude, ∼ 600 *µ*V; typical mean axonal spike amplitude, 4–15 *µ*V. The inset shows the distribution of branch lengths in the dataset (431 branches from 248 neurons across 14 networks; coefficient of variation = 0.386).

For each identified axonal branch, the time of arrival (TOA) at a given electrode was defined as the latency of spike detection relative to the soma. Conduction velocities at the start and end of the branch were estimated from TOA–distance relationships fitted by ordinary least squares to the first and last 20% of sampled points with a minimum of five data points contributing to each segment fit. The end-to-start velocity ratio, *ρ* = *v*_end_*/v*_start_, was required to satisfy a broad plausibility range, 0 < *ρ* < 2.5, chosen on the basis of velocity statistics from our laboratory’s HD-MEA dataset (mean 0.5 m/s, SD 0.12 m/s; 560 neurons, 9557 segments), consistent with published HD-MEA measurements [17, 31, 32]. No branch in the present dataset approached this bound, so the criterion was non-restrictive.

Branches were retained only if both *v*_start_ and *v*_end_ fits achieved *R*^2^ ≥ 0.95. After screening, the dataset consisted of 431 branches (from an initial 477) of length *L* = 600–4570 *µ*m, recorded from 248 neurons across 14 networks. Branches excluded by the *R*^2^ ≥ 0.95 criterion (minimum *R*^2^ ≈ 0.85) had a mean slowdown ratio of *ρ* ≈ 0.61, with mean fit qualities of 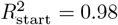 and 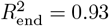, indicating that exclusion reflected noisier TOA–distance relations rather than a systematic bias in slowdown. All statistical analyses were performed in *Mathematica* and *Python* using custom scripts.

Before analyzing the statistical form of *ρ*, we asked whether its variability might reflect differences in axonal length (which correlates strongly with the number of branch points; *r* > 0.88 [29]). Across all 431 branches, *ρ* showed no detectable relationship with branch length (Pearson *r* = 0.018, *p* = 0.71; Spearman *r* = 0.044, *p* = 0.36). Thus, length variation does not account for the spread of *ρ*.

Four examples of TOA–distance relationships are shown in Figure 2, along with linearly fitted functions for estimation of end-to-start velocities, *ρ* = *v*_end_*/v*_start_. Log-log insets emphasize the continuous nature of the TOA–distance relationship. In all four exemplar cases of Figure 2, a slowdown phenomenon (*ρ* < 1) is clearly visible.

**FIG. 2.**
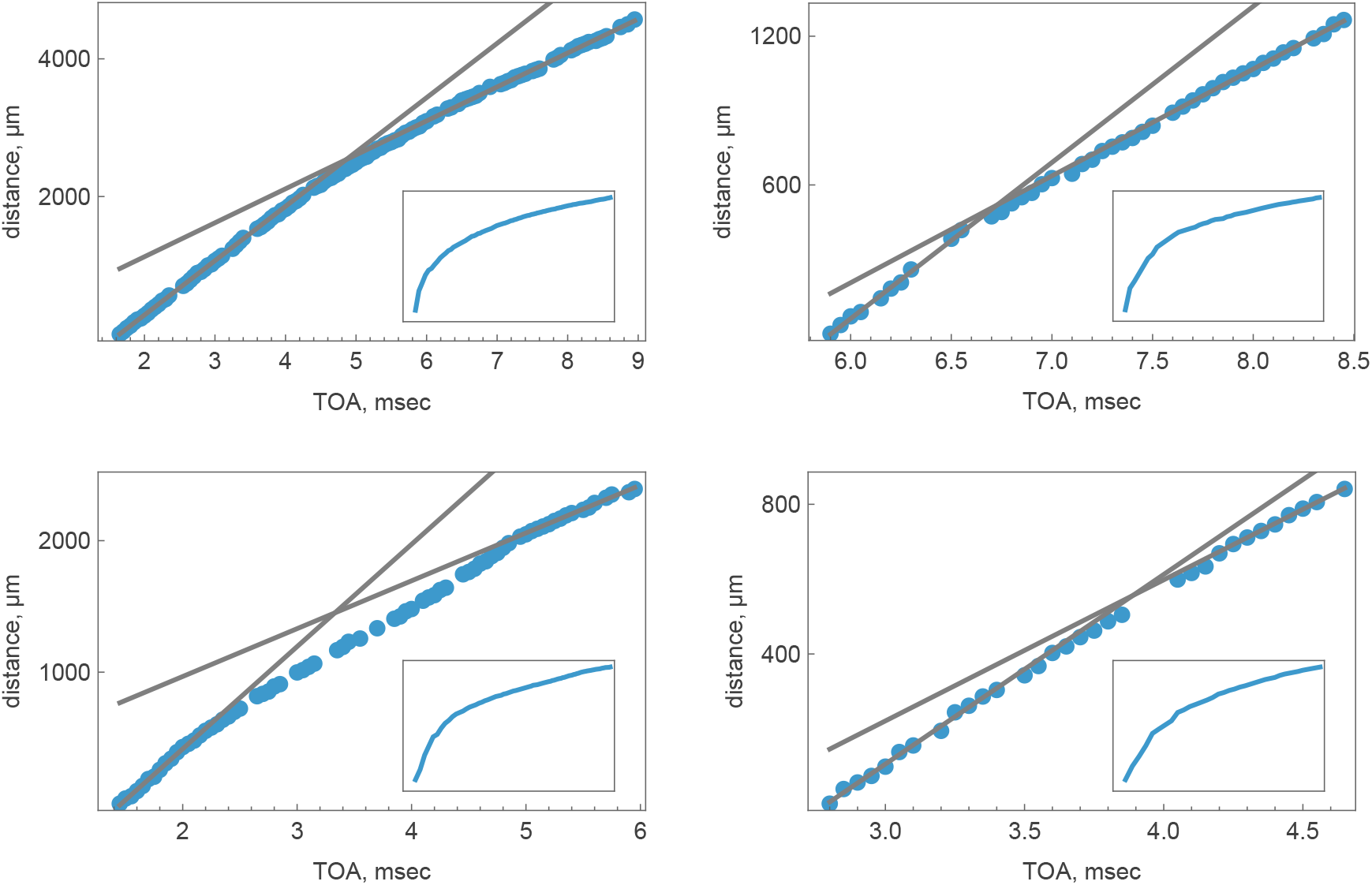
Time-of-arrival (TOA) versus distance for representative axonal branches, with linear fits to the initial and terminal segments used to estimate start and end conduction velocities, *v*_start_ and *v*_end_. Insets show the same data on log-log axes, emphasizing the smooth, continuous nature of the TOA–distance relationships. The two bottom panels correspond to segments from the neuron of Figure 1.

The distribution of *ρ* across 431 branches (Figure 3) is skewed, with typical branches slow by ∼ 30% (mean *ρ* ≈ 0.70, median *ρ* ≈ 0.68), and a small right tail for rare acceleration (approximately 6% of branches exhibiting *ρ* > 1). The coefficient of variation is 0.274, significantly less compared to the coefficient of variation of branch length in the same dataset, as shown in Figure 1 (ΔCV = 0.11, 95% CI [0.06, 0.16]). Were *ρ* merely a geometric reflection of length variability, its spread would be expected to equal or exceed that of length. The reduced variability of *ρ* therefore points to a normalization process in the axonal conduction machinery that compensates for geometric heterogeneity.

**FIG. 3.**
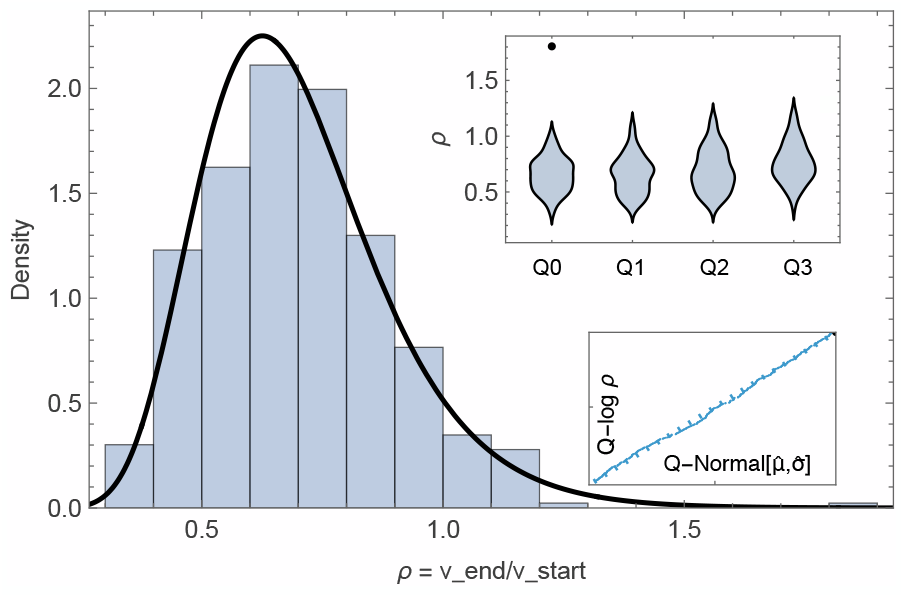
Distribution of conduction velocity ratios, *ρ* = *v*_end_*/v*_start_, across 431 axonal branches (coefficient of variation 0.27). The solid line shows the maximum-likelihood log-normal fit (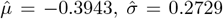 median *ρ* ≈ 0.68), representative of proportional compounding models. *Bottom-right inset* : Quantile–quantile (Q–Q) plot comparing ordered empirical log *ρ* values with expected quantiles of 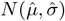; the near-linear alignment indicates that log *ρ* closely follows a normal distribution, i.e. *ρ* is log-normally distributed. *Top-right inset* : Violin representation of *ρ* stratified by quartiles of branch initial spike time (proxy for distance from soma). The preserved right-skewed shape across quartiles demonstrates distributional invariance; note the slight shift of the median *ρ* in distal (higher quartiles) branches. The black point at *Q*0 reflects a single outlier observation where the calculated initial velocity was 0.35 m/s, 2.4 SD units from the average initial velocity (0.65 m/s; *n* = 431 branches).

A maximum-likelihood estimation of the distribution shown in Figure 3 yields log 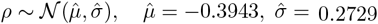, implying median 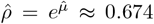 and mean 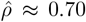, closely matching empirical values. 95% bootstrap confidence intervals (CI_95_) were estimated for the log-normal parameters (*µ, σ*) and the median of *ρ* using 2000 resamples, yielding *µ* ∈ [ −0.42, −0.37], *σ* ∈ [0.26, 0.29], and median *ρ* ∈ [0.66, 0.69]. The Q–Q plot for log *ρ* (inset to Figure 3) is near-linear with minor deviations.

To verify that the observed log-normal form of slowdown is not an artifact of model choice, we performed a series of robustness checks. To encompass the main mechanistic possibilities, we examined three representative families, the *Gamma, log-normal*, and *log-logistic* distributions, providing a minimal yet comprehensive basis for assessing which process best accounts for the observed form of *ρ*. Information criteria favored the log-normal model (AIC = −232.18; BIC = −224.05), with the Gamma distribution close behind (ΔAIC = ΔBIC = 1.31) and therefore statistically indistinguishable. However, the Gamma family represents an additive generative process, in which variability accumulates through independent increments. This is less physiologically well grounded: excitability mechanisms, particularly sodium conductance, gating kinetics, and local impedance interactions, operate through proportional effects on current and voltage, not through additive time delays. The log-logistic model performed poorly (ΔAIC = ΔBIC = 12.86), providing no credible support for a regulated-multiplicative process. For completeness, we also fitted an *extreme-value* (Gumbel) distribution, which yielded ΔAIC = ΔBIC = 6.36 relative to the log-normal. Note that an extreme-value process is mechanistically difficult to reconcile with the data: The smooth convexity of time-of-arrival relations, the independence of *ρ* from branch length, and the near-normal distribution of log *ρ* all indicate that slowdown arises from the compounded effect of many small influences rather than from a bottleneck along the axon.

To examine whether hierarchical dependencies among branches, neurons, and networks influence the observed distribution of conduction slowdown, we aggregated *ρ* values at successive levels and re-fitted the log-normal model to each aggregate. At the neuron level, branch-specific *ρ* values were averaged to yield one representative value per neuron, and at the network level, neuron means were further averaged to produce one value per network. The fitted parameters remained stable across levels: Branch level (reported above): 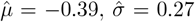; Neuron level: 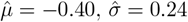 Network level: 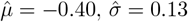). The corresponding medians (≈ 0.67) and means ( ≈ 0.69) were nearly invariant. The constancy of *µ* and the median indicates that the central tendency of slowdown is invariant to aggregation. The mild reduction of *σ* with aggregation reflects within-neuron and within-network correlations of modest magnitude. Thus, clustering exists but does not distort the log-normal form nor the overall 30% slowdown ratio.

Stratifying *ρ* by branch initial spike time (Figure 3, top-right inset), a proxy for branch position relative to the soma, yielded overlapping distributions with stable shape: 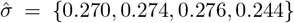 and median *ρ* = {0.648, 0.640, 0.670, 0.743 for Q0–Q3}, respectively. The slight increase in *ρ* for branches that begin more distally is expected: conduction velocity is already lower at the onset of distal branches, reflecting the progressive deceleration evident in individual trajectories (see, for example, the left panel of Figure 4).

**FIG. 4.**
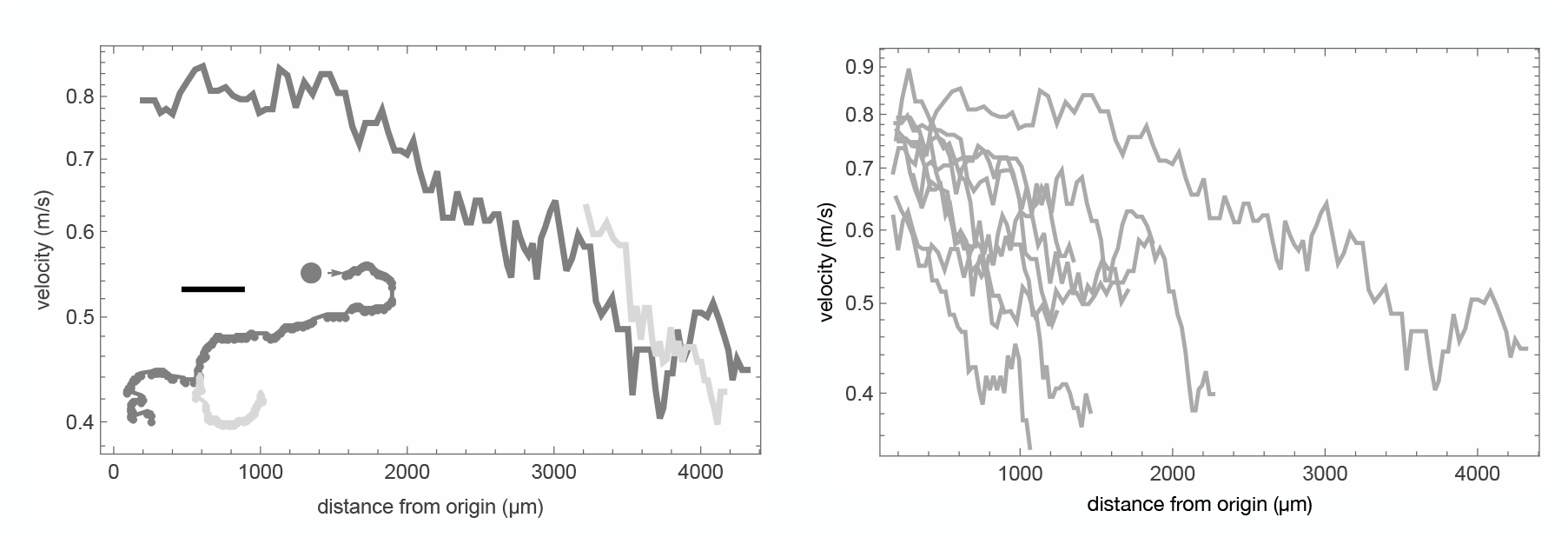
*Left* : Log-linear conduction-velocity profile along an individual bifurcating axon (moving average ≈ 350, *µ*m). The time delay between the triggering spike (its location depicted by a gray-filled circle) and the first point of the long branch is 250 *µ*s. Grayscale corresponds to the branch colors shown in the lower-left inset; each branch inherits its initial velocity from the terminal velocity of its parent. Scale bar: 500, *µ*m. *Right* : Log-linear conduction-velocity profiles of axons with different lengths, all originating near the soma.

As an additional diagnostic, we examined the ratio between the average and initial conduction velocities for each branch, denoted 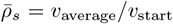, and the ratio between the end and average conduction velocities, denoted 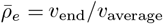. These measures provide a complementary view to the end-to-start ratio (*ρ* = *v*_end_*/v*_start_), summarizing the overall slowing along the trajectory. Across all branches, 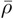 exceeded 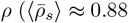 and 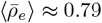 vs. 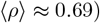. The distributions of 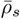 and 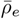 were likewise right-skewed, with approximately 18% and 10% of branches, respectively, exhibiting values greater than unity. These findings suggest that the overall deceleration is gradual and systematic. The data shown in Figure 4 supports this idea of monotonic, gradual deceleration. The left panel shows a log-linear conduction-velocity profile along one bifurcating axon (moving average ≈ 350, *µ*m). Note the gradual decline and how the branch inherits its initial velocity from the terminal velocity of its parent. Several log-linear conduction-velocity profiles of axons with different lengths, all originating near the soma, are shown in the right panel of Figure 4. The slowdown is progressive and monotonic rather than step-like.

Moreover, as shown in the left panel of Figure 5, histograms of *v*_*start*_ (blue) and *v*_*end*_ for 200 axons with an even more strict initial time of arrival (TOA), ≤ 150 *µ*s relative to the triggering somatic spike, indicate that for *v*_*start*_, the average velocity (0.66 m/s) translates to branch initiation at ≤ 100 *µ*m from the triggering spike. The normal model fits the data exceptionally well: the goodness-of-fit *p*-value is 0.922, the AIC is −254.26, and the sample skewness is nearly zero (0.101), all indicating a symmetric distribution consistent with normality. The log-normal alternative performs worse on every metric, yielding a lower *p*-value (0.518), a higher AIC (−245.71), and more pronounced skewness in the log-domain ( −0.541). In contrast, the *v*_*end*_ dataset exhibits the opposite pattern. Here, the log-normal model is favored, with a high *p*-value (0.864), the lowest AIC ( −354.50), and near-zero skewness after log transformation ( −0.177). The normal model, by comparison, produces a lower *p*-value (0.388), a substantially higher AIC (349.13), and a noticeably right-skewed distribution (skewness = 0.358). Taken together, these results indicate a clear transition along the axonal trajectory, from an approximately normal distribution at the start to a log-normal distribution at the end. Here too, there is practically no correlation between *ρ* and axonal length for the 200 axons whose histograms are depicted in the left panel. The scatter is effectively random, indicating no statistically significant correlation between axonal length and slowdown ratio. Shown are the linear fit and 95% confidence interval. Pearson correlation: *r* = 0.05, *p* = 0.48; Spearman rank correlation: *r*_*s*_ = 0.08, *p* = 0.26.

**FIG. 5.**
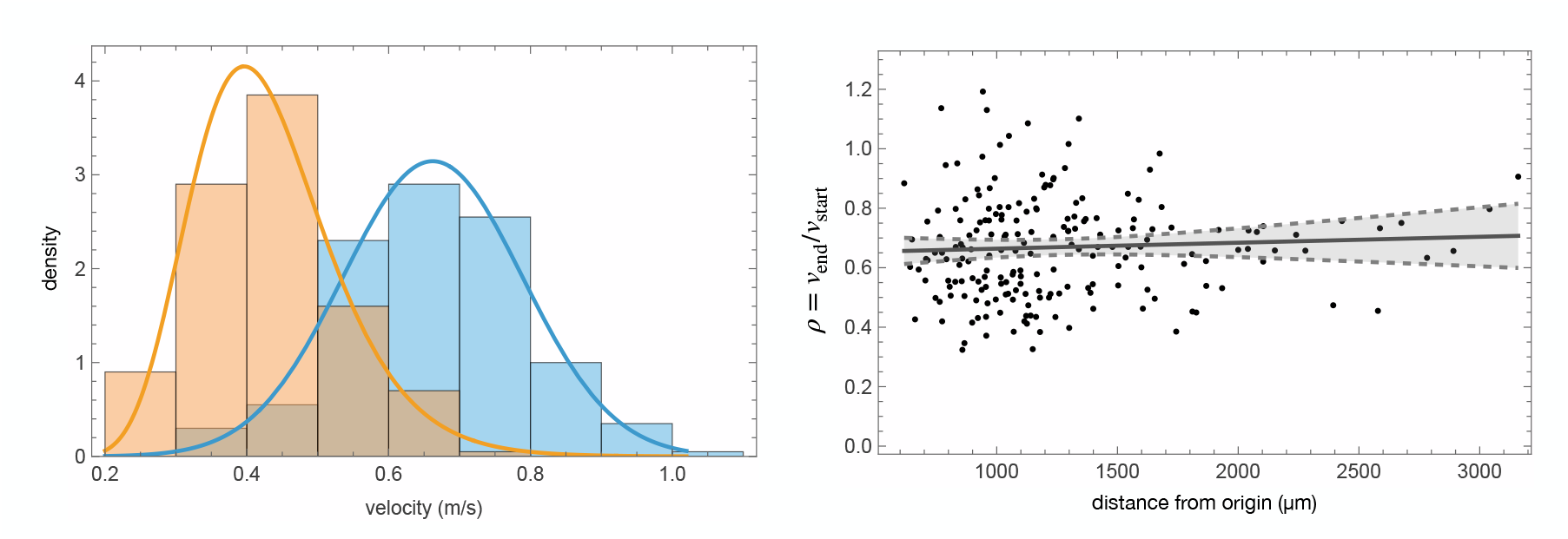
*Left* : Histograms of *v*_*start*_ (blue) and *v*_*end*_ are shown for 200 axons with a strict initial time of arrival (TOA), ≤ 150 *µ*s relative to the triggering somatic spike. For *v*_*start*_, the average velocity (0.66 m/s) translates to branch initiation at ≤ 100 *µ*m from the triggering spike. The normal model fits the data exceptionally well: the goodness-of-fit *p*-value is 0.922, the AIC is 254.26, and the sample skewness is nearly zero (0.101), all indicating a symmetric distribution consistent with normality. The log-normal alternative performs worse on every metric, yielding a lower *p*-value (0.518), a higher AIC ( −245.71), and more pronounced skewness in the log-domain ( −0.541). In contrast, the *v*_*end*_ dataset exhibits the opposite pattern. Here, the log-normal model is favored, with a high *p*-value (0.864), the lowest AIC ( −354.50), and near-zero skewness after log transformation ( −0.177). The normal model, by comparison, produces a lower *p*-value (0.388), a substantially higher AIC (349.13), and a noticeably right-skewed distribution (skewness = 0.358). Taken together, these results indicate a clear transition along the axonal trajectory, from an approximately normal distribution at the start to a log-normal distribution at the end. *Right* : *ρ* as a function of axonal length for the 200 axons whose histograms are depicted in the left panel. The scatter is effectively random, indicating no statistically significant correlation between branch length and slowdown ratio. Shown are the linear fit and 95% confidence interval. Pearson correlation: *r* = 0.05, *p* = 0.48; Spearman rank correlation: *r*_*s*_ = 0.08, *p* = 0.26.

Having established these empirical regularities, we turn to their mechanistic and physiological implications. The observed approximate invariance of conduction slowdown across branch lengths and aggregation levels is consistent with propagation speed being governed by a stochastic process in which local influences compound to produce a distribution that remains stable across branches, neurons, and networks. The essential empirical constraint is the invariance itself: any viable mechanistic account must reproduce the stability of the distribution under structural diversity. This behavior is consistent with, though not exclusive to, multiplicative dynamics, in which each axonal segment rescales velocity by a fractional factor reflecting its geometry or biophysical state, with the cumulative effect yielding the observed statistical form.

Processes of this sort are familiar from stochastic transport in disordered media, where numerous small influences combine to generate log-normal or related right-skewed distributions. In axons, several mechanisms could instantiate such invariance: current splitting at bifurcations, impedance mismatches, gradual diameter taper, and state-dependent channel kinetics. None of these mechanisms is mutually exclusive; together they may constitute the effective stochastic law governing propagation.

A particularly relevant example for one such mechanism is the inactivation of voltage-gated sodium channels, whose fractional availability is activity-dependent over wide temporal scales [4, 33]. Because channel gating enters multiplicatively into total sodium conductance, variations in availability modulate conductance proportionally. Since conduction velocity scales with instantaneous sodium conductance, such proportional modulation can compensate for structural variability and support robust function.

From a physiological standpoint, the robustness of conduction to structural variability is fundamental. Neural tissue operates within a geometrically irregular and dynamically mutable medium, where precise architectural reproducibility is neither attainable nor required. That axonal propagation remains statistically invariant despite such variability implies that the organization of excitability is tuned for functional reliability rather than for precision of structure. Proportional or self-normalizing regulation ensures that local differences in geometry or conductance compound without degrading transmission fidelity. Conduction dynamics are stabilized by the same scale-invariant statistical organization that sustains excitability itself [13], rather than by structural scaling, enabling faithful signal transmission in a structurally heterogeneous substrate.

The authors thank Tamar Galateanu and Leonid Odessky for technical assistance, and Erez Braun, Naama Brenner, Hana Salman, and Noam Ziv for insightful discussions. This work was partially supported by the Schaefer Scholars Program at Columbia University’s Vagelos College of Physicians and Surgeons.

